# A Cyclic Phosphoramidate Prodrug of 2’-deoxy-2’-fluoro-2’-C-methylguanosine for the Treatment of Dengue Infection

**DOI:** 10.1101/2020.04.13.040287

**Authors:** Ratna Karuna, Fumiaki Yokokawa, Keshi Wang, Jin Zhang, Haoying Xu, Gang Wang, Mei Ding, Wai Ling Chan, Nahdiyah Abdul Ghafar, Andrea Leonardi, Cheah Chen Seh, Peck Gee Seah, Wei Liu, Rao PS Srinivasa, Siew Pheng Lim, Suresh B Lakshminarayana, Ellie Growcott, Sreehari Babu, Martijn Fenaux, Weidong Zhong, Feng Gu, Pei-Yong Shi, Francesca Blasco, Yen-Liang Chen

**Affiliations:** Novartis Institute for Tropical Diseases, Singapore; Novartis Institutes for BioMedical Research, Emeryville, CA, USA; Novartis Institutes for BioMedical Research, East Hanover, NJ, USA; Novartis Global Drug Development, Shanghai, China; Novartis Institutes for BioMedical Research, Cambridge, MA, USA

**Keywords:** dengue, nucleotide analog, monophosphate prodrug, cyclic phosphoramidate, nucleoside triphosphate, polymerase inhibitor

## Abstract

Monophosphate prodrug analogs of 2’-deoxy-2’-fluoro-2’-*C*-methylguanosine have been reported as potent inhibitors of hepatitis C virus (HCV) RNA-dependent RNA polymerase. These prodrugs also display potent anti-dengue activities in cellular assays although their prodrug moieties were designed to produce high levels of triphosphate in the liver. Since peripheral blood mononuclear cells (PBMCs) are one of the major targets of dengue virus, different prodrug moieties were designed to effectively deliver 2’-deoxy-2’-fluoro-2’-*C*-methylguanosine monophosphate prodrugs and their corresponding triphosphates into PBMCs after oral administration. We identified a cyclic phosphoramidate prodrug **17** demonstrating a well-balanced anti-dengue cellular activity and *in vitro* stability profiles. In dogs, oral administration of **17** resulted in high PBMC triphosphate level, exceeding TP_50_ (the intracellular triphosphate concentration at which 50% of virus replication is inhibited) at 10 mg/kg. Compound **17** demonstrated 1.6- and 2.2 log viremia reduction in the dengue mouse model at 100 and 300 mg/kg twice daily, respectively. At 100 mg/kg twice daily, the terminal triphosphate concentration in PBMCs reached above TP_50_, defining for the first time the minimum efficacious dose for a nucleos(t)ide prodrug. In the two-week dog toxicity studies at 30 to 300 mg/kg/day, no observed adverse effect level (NOAEL) could not be achieved due to pulmonary inflammation and hemorrhage. The preclinical safety results suspended further development of **17**. Nevertheless, present work has proven the concept that an efficacious monophosphate nucleoside prodrug could be developed for the potential treatment of dengue infection.

## INTRODUCTION

The mosquito-borne dengue virus is endemic to tropical and sub-tropical regions throughout the world, making dengue fever the most important mosquito-borne viral disease afflicting humans. Its global distribution is comparable to that of malaria, with an estimated 2.5 billion people at risk for epidemic transmission (1). There has been steady increases in countries affected and incidence since the 1950s and recent estimates suggest annual rates of 390 million cases accompanied by 20,000 deaths (2).

Dengue viruses (DENVs) can be further classified into four different serotypes (DENV-1 to −4), all of which can lead to disease symptoms with varying severity. Secondary infection by a different serotype may increase the risk of severe dengue diseases. While diagnosis of dengue infection can be rapid and simple, serotype distinction requires additional instrumentation, usually in a laboratory setting. Thus, the ideal treatment for dengue fever should possess pan-serotype activities (3). Recently, a dengue vaccine was approved in certain countries but is recommended only for individuals with prior DENV exposure. This limits its use as well as necessitating pre-vaccination screening (4). No antivirals are currently available for the treatment of dengue.

DENV is an enveloped, positive strand RNA virus belonging to the *Flaviviridae* family and the genus *Flavivirus*. Medically important viruses in this class include Yellow Fever virus (YFV), Japanese encephalitis virus (JEV), West Nile virus (WNV), and Zika virus (ZIKV). The dengue viral genome encodes three structural (C-prM-E) and seven nonstructural (NS1-NS2A-NS2B-NS3-NS4A-NS4B-NS5) proteins. The nonstructural protein NS5 contains both methyltransferase and RNA-dependent RNA polymerase (RdRp) activities. The RdRp is a viral specific enzyme which catalyzes the replication of viral RNA from its own complementary template. It is essential for viral replication and an attractive target for therapeutic intervention (5).

Nucleoside/Nucleotide analogs are a highly successful compound class of antivirals, as exemplified in the treatment of human immunodeficiency virus (HIV), herpes simplex virus (HSV), hepatitis B virus (HBV), and hepatitis C virus (HCV) (6). These inhibitors are converted to their active nucleoside triphosphate forms by host-cell machinery and inhibit the synthesis of viral RNAs or DNAs by acting as ‘chain terminators’ or substrate mimics (7). Because the modified nucleoside triphosphates must be recognized by the highly conserved active site of DENV polymerase, they have a high likelihood of pan-DENV serotype activity and a high barrier to drug resistance (8). These features make them very attractive for dengue drug development (9). We previously reported an adenosine-based nucleoside (Fig. 1A, NITD-008 (**1**), 7-deaza-2’-*C*-ethynyl-adenosine) which potently inhibited DENV both *in vitro* and *in vivo*. However, the development of NITD-008 was terminated due to insufficient safety profile (10). 4’-azido-cytidine (R-1479 (**2**)) and its ester prodrug, balapiravir (**3**) (Fig. 1A), were originally developed for the treatment of HCV, but their development were terminated due to hematologic adverse events such as lymphopenia (11). Azido-cytidine (**2)**was weakly active against dengue (12) and Balapiravir (**3**) failed to reduce viral load in dengue patients (13).

**FIG 1.**
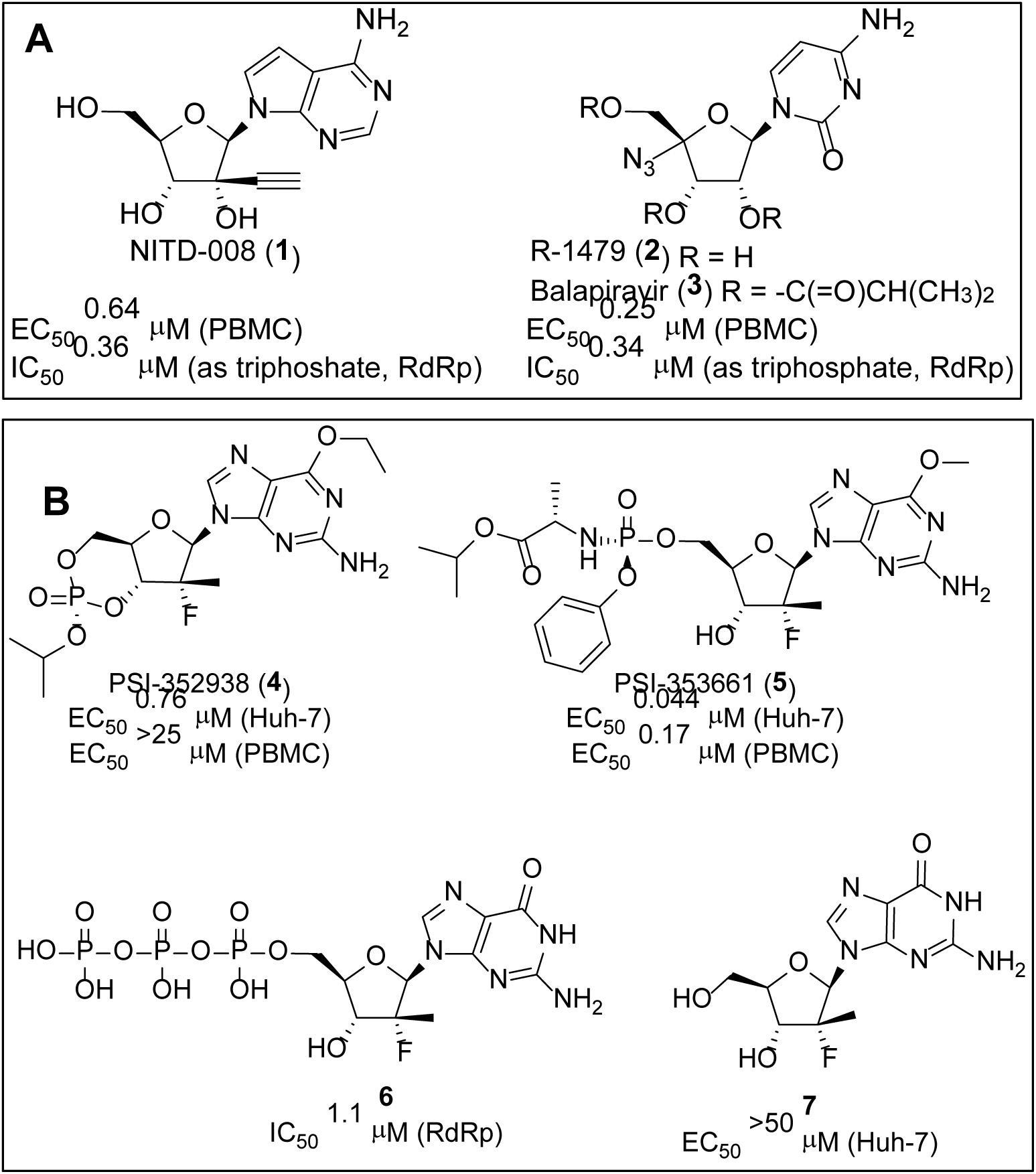
**(A)** Structures and *in vitro* biological profile of NITD-008 **1**, R-1479 **2**, and balapiravir (**3**). **(B)** Structures and *in vitro* biological profile of PSI-352938 **4**, PSI-353661 **5**, and their corresponding triphosphate **6** and nucleoside **7** metabolites

It is not uncommon that many nucleoside analogs suffer the lack of biological activities in cellular assays due to poor intracellular conversion to their triphosphates. In particular, conversion of nucleoside analogs into their nucleotide or nucleoside monophosphate is often rate-limiting or non-productive (14). On the other hand, the unprotected monophosphate species are poor drug candidates as they have inadequate cellular permeability due to the inability of negatively charged phosphates to cross the cell membrane. To circumvent the problem, the monophosphate prodrug approach has been developed to deliver the nucleoside monophosphate directly into target cells (15). This prodrug approach has proven to be effective in improving the therapeutic potential of antiviral and anticancer nucleosides (16). For instance, the uridine-based monophosphate prodrug sofosbuvir is a key component in a number of HCV combination therapies (17). The excellent efficacy of sofosbuvir is due to the efficient delivery of the nucleoside triphosphate into the target organ, liver (18).

Similarly, prodrugging nucleotides to deliver a high level of active triphosphates into PBMCs as has been successfully demonstrated in tenofovir and GS-5734 (19) alafenamide (20) for the treatment of HIV and Ebola virus, respectively. Due to the ester moiety, rapid elimination of the prodrug was observed. However, GS-5734 was also rapidly distributed into PBMCs and the corresponding triphosphate level reached maximum of 2 hours in PBMC (19). The prodrug tenofovir alafenamide was only transiently present in plasma (T_1/2_ ∼30 min) when dosed orally to dogs. However, the exposure was sufficient to drive a high and sustained level of the active metabolite in PBMCs (20).

In 2010, Pharmasset (acquired by Gilead in 2012) reported two nucleotide prodrugs of 2’-deoxy-2’-fluoro-2’-*C*-methylguanosine, PSI-352938 (**4**) (21, 22) and PSI-353661 (**5**) (23) as potent inhibitors of HCV replication (Fig. 1A). Their common guanosine-based nucleoside triphosphate **6** was reported to be a potent inhibitor of HCV NS5B polymerase with IC_50_ value of 5.94 µM. We evaluated these prodrugs and the active triphosphate in our PBMC dengue plaque and dengue RdRp enzyme assays, respectively. We subsequently embarked on the optimization of the prodrug moieties to deliver the active triphosphate **6** into peripheral blood mononuclear cells (PBMCs), one of the major dengue replication sites (24).

Herein we report our research leading to a cyclic phosphoramidate prodrug of 2’-deoxy-2’-fluoro-2’-*C*-methylguanosine for the treatment of dengue starting from the liver-targeting prodrugs. The optimized prodrug **17** resulted in high triphosphate loading in PBMCs after oral administration in dogs. In addition, compound **17** demonstrated oral efficacies at 100 mg/kg twice daily in the dengue mouse model with high triphosphate concentration in PBMCs, while the liver-targeting prodrug **5** failed to reduce viremia at an even higher dose. Based on the correlation of the triphosphate levels in PBMCs and *in vivo* efficacy, we defined the minimum efficacious dose for a nucleoside/nucleotide prodrug. We also described the *in vitro* and *in vivo* preclinical characterization of **17** as well as the development work towards safety assessment.

## MATERIALS AND METHODS

### Material

All nucleoside/nucleotide compounds were synthesized at Novartis Institute for Tropical Diseases (NITD) as described in the main text. The solid dispersion batch of compound **17** was manufactured by the Chemical and Pharmaceutical Profiling (CPP) unit of Novartis in Shanghai, China. The solid dispersion formulation consisted of 20% (w/w) active ingredient, 40% (w/w) hypromellose acetyl succinate (HPMC-ASLF, Shin-Etsu Chemical, Tokyo, Japan), 35% (w/w), hypromellose (HPMC-E3, Shin-Etsu Chemical, Tokyo, Japan) and 5% (w/w) sodium lauryl sulfate (SLS, Sigma-Aldrich, St. Louis, MO). The analytical standard for triphosphate measurement, 8-Bromoadenosine 5′-triphosphate (Br-ATP), and ion-pairing reagent hexylamine were from Sigma-Aldrich. Acetonitrile and ammonium acetate used for LC-MS/MS mobile phases were from Merck (Darmstadt, Germany). All other solvents, reagents, and chemicals were either of molecular biology grade or of the highest chemical grade available from Sigma-Aldrich or Thermo Fisher Scientific (Waltham, MA) unless otherwise mentioned.

Different species of pooled liver S9 fraction (Gentest™, Corning, NY) and intestinal S9 fraction (XenoTech, Kansas City, KS) were of mixed gender for human and male for all other species. Co-factor NADPH is from Sigma-Aldrich (St. Louis, MO). Different species of plasma were all mixed genders and obtained from Seralab (West Sussex, UK). Cryopreserved human PBMCs (individual donors) were purchased from AllCells (Alameda, CA) or ReachBio (Seattle, WA). Written consent from the donors was available for all samples. All experiments involving human matrices were approved by the Institutional Review Board of Novartis prior to the start of the experiments. PBMCs from other species (pooled) were from 3H Biomedical (Uppsala, Sweden). C6/36, THP-1, KU812, K562, 293T and BHK-21 were from American Type Culture Collection (ATCC, Manassas, VA).

Vacutainer® CPT™ (Cell Preparation Tube with sodium citrate, 4 ml draw capacity) was from BD Biosciences (Franklin Lakes, NJ). The collagen I coated plates were from Thermo Fisher Scientific. HEPES buffer, RPMI 1640 medium, penicillin-streptomycin were from Life Technologies (Carlsbad, CA). PhosSTOP (phosphatase inhibitor) and protease inhibitor cocktail (cOmplete™) tablets were from Roche Applied Science (Penzberg, Germany).

### Stability in plasma, liver and intestinal S9

Frozen pooled plasma (K_3_EDTA) was thawed from −20°C and centrifuged at 2643 × *g* for 5 minutes (min) at ambient temperature and any hemin plug was discarded. Plasma was then diluted in Dulbecco’s phosphate buffer saline (PBS) to achieve 50% concentrated solution with nominal pH of ∼7.4±1. A pre-warmed diluted plasma and compound in methanol (1 µM final concentration) was briefly vortexed at time point zero. Incubation was performed in a shaking water bath (37°C) and subsequent time points were taken (5, 15, 30, 60, 120 min). Samples were quenched with 4 volumes of ice-cold acetonitrile (containing internal standard), centrifuged, and supernatant analyzed by LC-MS/MS. Half-life ((t_1/2_) was then calculated from the parent depletion.

Frozen pooled liver or intestinal S9 fraction was thawed from −20°C and diluted in PBS with NADPH (1 mM final concentration) to 2 mg/ml. The PBS was pre-warmed at 37°C for 10 min. The reaction was initiated by addition of compound in methanol (1 µM final concentration). The sample plates were incubated on a shaker (37°C). Sequential samples were removed at designated time points (0, 5, 15, 30, 60, 120 min) and quenched with 4 volumes of ice-cold acetonitrile (containing internal standard), centrifuged and supernatant reconstituted in water (acetonitrile:water, 1:1 v/v). Samples were then analyzed by LC-MS/MS and half-life (t_1/2_) was obtained.

For all *in vitro* stability assays, a generic LC-MS/MS method was used to assess parent depletion. Briefly, separation was performed on a 50 × 2 mm, 4 micron, Synergy Polar-RP column (Phenomenex, Torrance, CA) using a fast gradient elution of 400 µl per minute (5% to 95% B in 0.8 min and kept at 95% B for another 1.6 min). Mobile phase was 0.1% formic acid in water (A) or acetonitrile (B). Detection was performed using TSQ Quantum™ Discovery Max (Thermo Fisher Scientific) with electrospray ionization (ESI) in positive mode. Stability was determined semi-quantitatively from the peak area ratios of analyte:internal standard (diazepam) and half-life (t_1/2_) was calculated based on the rate of compound depletion.

### *In vitro* antiviral assays

Antiviral assays in PBMCs were performed as previously described (12). Briefly, cryopreserved PBMCs were thawed according to the manufacturer’s instruction and re-suspended in RPMI medium supplemented with 1% penicillin-streptomycin solution. The DENV was pre-incubated with 0.38 µg/ml chimeric 4G2 antibody for 30 min at 4°C to form a virus-antibody complex before it was added to the PBMCs at multiplicity of infection (MOI) of 1. The plate was further incubated at 37°C for 1 hour before addition of serially diluted compounds. The viral titers in the culture fluids were quantified using a plaque assay at 48 hour post-infection. EC_50_ was calculated by Prism (GraphPad Software, La Jolla, CA) using the equation for a sigmoidal dose-response (variable slope).

Huh-7 dengue replicon assay was previously described (25). For 293T dengue plaque assay, 3 × 10^4^ cells were seeded in a 96-well collagen I coated plate in 100 µl of media (DMEM supplemented with 10% fetal bovine serum and 1% penicillin-streptomycin) one day prior to infection. The media was removed and the cells were infected with the virus-antibody complex with MOI of 3 in the same media without serum for 1 hour at 37°C. The media was then removed and replaced with DMEM media supplemented with 2% fetal bovine serum and 1% penicillin-streptomycin for another 2 days. The EC_50_ was calculated using a plaque assay as described before. THP-1 and KU812 assays were performed as described in the publications (26). Cytotoxicity assay was also performed as previously described (27). K562 assay was performed similarly with the MOI of 1.

RdRp assay was conducted as previously described (12). Briefly, a 244-nucleotide RNA with the sequence 5’-(TCAG)_20_(TCCAAG)_14_(TCAG)_20_-3’ was used as a template (28). Compounds with various concentrations were mixed with the RNA template (100 nM), dengue RdRp (100 nM), 0.5 µM of BBT-GTP and 2 µM of ATP, CTP and UTP in the buffer containing 50 mM Tris HCl pH 7.5, 10 mM KCl, 0.5 mM MnCl_2_ and 0.01% Triton X-100 for 120 minutes. The amount of substrate produced and IC_50_ was determined as described in the publication (29).

### *In vitro* triphosphate conversion studies in PBMCs

Cryopreserved PBMCs were thawed and incubated in RPMI medium containing 2% fetal bovine serum, 1% penicillin-streptomycin, and a designated concentration of the prodrug. After the intracellular conversion into corresponding triphosphate reached steady state (24 hours at 37°C in the incubator), the cells were spun down for 10 min (ambient temperature, 135 × *g*) and washed with cold 0.9% NaCl solution in 1 mM HEPES. The wash buffer was carefully removed with a micropipette and cell lysis carried out before compound measurement.

For investigating the triphosphate conversion kinetic, a sequential time points were taken (0, 3, 7, 24, 48 hours) upon prodrug incubation in human PBMCs. For half-life experiment, compound **12** (100 µM) was incubated in human PBMCs for 24 hours, the media was replaced with fresh media without compounds and time points were subsequently taken (0, 2, 4, 8, 24, 32, 48 hours).

### TP_50_ determination

TP_50_ is defined as the intracellular triphosphate concentration at which 50% of the virus replication is inhibited. Due to analytical sensitivity, direct measurement of the triphosphate concentration at the EC_50_ of the prodrug was challenging. Hence, the TP_50_ was derived using Michaelis-Menten kinetic with increasing prodrug concentrations in human PBMCs (12). Briefly, cryopreserved human PBMCs were incubated with compound **17** (3, 10, 30 100 µM). The cells were prepared and lysed and the PBMC triphosphate was analyzed by LC-MS/MS as described in other parts of this Method section. The intracellular triphosphate concentrations were plotted against the prodrug concentrations used in the incubation. TP_50_ was derived using Michaelis-Menten kinetic with the formula of *Y = A[X]/B + [X]*, where *Y* is the triphosphate concentration, *A* is the calculated maximum triphosphate concentration extrapolated from the graph, *[X]* is the prodrug concentration, and *B* is the calculated prodrug concentration at which its triphosphate reached half of its maximal value.

### PBMC lysis

Cell lysis buffer containing 50 mM Tris-HCl at pH 7.5, 150 mM NaCl, 1% IGEPAL® CA-630, 1 mM phenylmethane sulfonyl fluoride (PMSF), 1 × protease inhibitor stock solution (Roche), and 1 PhosSTOP tablet was freshly made in 10 ml solution and used within 2 hours. The 50 × protease inhibitor stock solution was made by dissolving one protease inhibitor cocktail tablet in 1 ml water. For rodent PBMCs, the concentration of protease inhibitor was increased five-fold to ensure the stability of the triphosphate in the lysate.

PBMCs were lysed by adding cell lysis buffer at 10 million cells/ml and incubated at room temperature for 10 min. The cell debris was then spun down at 15,800 × *g* for 20 min (4°C). The lysate was transferred into new tubes, snapped frozen in liquid nitrogen and stored at −80°C until triphosphate LC-MS/MS analysis.

### PBMC isolation from *in vivo* studies

To increase sensitivity of triphosphate detection, PBMCs were lysed at the concentration of 30 million cells/ml from each dog or pooled AG129 mice. To obtain PBMCs from animals, blood was drawn from the animals to Vacutainer® CPT™ tubes at indicated time points. The CPT tubes were centrifuged at ambient temperature for 20 min (1500 × *g*). A whitish layer (PBMCs) under plasma layer was pipetted into a 15 ml conical centrifuge tube. PBS (10 ml) was added and the mixture was centrifuged at ambient temperature for another 10 min (700 × *g*). The plasma supernatant was then carefully aspirated using a vacuum pump while ensuring the PBMC pellet remained at the bottom of the tube. The pellet was then resuspended with PBS (1 ml) and vortexed at the lowest setting. The suspension was transferred to an Eppendorf tube and lysed as described above. The blood was processed immediately to ensure minimum degradation of the triphosphate. The samples (cell lysate) were stored or shipped in −80°C for triphosphate LC-MS/MS analysis.

### LC-MS/MS analysis

Methods with slight variation were used for the multiple studies conducted at different sites. The basic principles of all the methods used were similar and comparable results were obtained. The intact prodrugs and free nucleoside metabolite **7** in plasma were measured after protein precipitation followed by reversed-phase liquid chromatography (LC) and tandem mass spectrometry (MS/MS) with ESI in positive mode. The intracellular triphosphate **6** analysis was carried out after protein precipitation of the PBMC cell lysate, ion-pairing to retain the triphosphate on a reversed-phase LC column, and MS/MS with ESI in negative mode.

Analysis was most challenging in the rodent matrix due to the *ex-vivo* instability of the intact prodrug in plasma and triphosphate in cell lysate. The methods used were as follows. For the intact prodrug and metabolite **7**, an inhibitor cocktail of NaF and citric acid (20 mM NaF and 40 mM citric acid final concentration) was added to the blood collection tubes. Plasma was obtained by centrifuging the blood for 5 min (4°C) at 10,000 × *g*. 175 µl extraction solution mixture (acetonitrile:methanol:acetic acid, 90:10:0.2 v/v) containing a generic internal standard (warfarin) was added to 25 µl plasma. The sample plates were shaken and then centrifuged for 10 min (4°C, 2884 × *g*). The supernatant was collected and 5 µl was injected to the LC-MS/MS system. Mobile phase was 20 mM ammonium acetate + 1% acetic acid (A) or in 100% acetonitrile (B). Gradient elution was performed on a Hydro-RP column (100 × 3 mm, 2.5 micron, Phenomenex, Torrance, CA) at 600 µl/min: 0% to 80% B in 2 min and kept at 80% B for another 1.5 min. MS/MS detection was performed with 4000 QTRAP® (Sciex, Framingham, MA). MRM transition of 489.4/310.2 and 300.3/152.1 were used for prodrug **17** and metabolite **7**, respectively. Calibration and quality control (QC) samples used matched-matrix (e.g. naïve AG129 mouse plasma) and the lower limit of quantification (LLOQ) obtained was 15.75 nM for both analytes.

For PBMC triphosphate analysis, a higher concentration of protease inhibitor cocktails was used during cell lysis of rodent samples (see PBMC lysis section). An equal amount of acetonitrile:water mixture (1:1) containing internal standard (Br-ATP) was added to the cell lysate. The mixture was vortexed and centrifuged (1,431 × *g* at ambient temperature) for 10 min. The supernatant (5 µl) was then injected to the LC-MS/MS system (4000 QTRAP®, Sciex, Framingham, MA). Ion-pair chromatography was used for separation on a 5 × 2 mm, 3 micron, Gemini-NX column (Phenomenex, Torrance, CA). Mobile phase A and B was water and acetonitrile mixture, respectively, containing 5 mM ammonium acetate and 5 mM hexylamine, buffered at pH 8.5. Gradient elution (700 µl/min) was achieved with 5% to 60% mobile phase B within 3.5 min. Detection was in negative mode with MRM transition 538.2/159.0 and 585.9/159.0 for the triphosphate **6** and Br-ATP, respectively. Matched-matrix (e.g. mouse PBMC lysate) with similar cell concentration as the samples were used for calibration and QC. The LLOQ was 0.065 pmol/3 million cells.

The triphosphate concentration obtained from LC-MS/MS analysis was expressed as mol per number of cells. The triphosphate concentration per cell volume (in µM) was calculated using the corpuscular volume of 283 fL for PBMC (30).

### *In vivo* studies

All Novartis animal studies were approved by institutional review board at different sites and different authorities where the experiments were carried.

The dog pharmacokinetic studies of compound **12, 14, 17**, and **18** were conducted using beagle dogs by intravenous administration (i.v., 0.5 mg/kg) or oral gavage (p.o., 3 mg/kg for each compound and 15 mg/kg only for **(17**). A solution formulation containing 20% PEG300, 5% Solutol HS-15, and 5% dextran in water was used for both the i.v. and p.o. doses. Blood samples (3.5 ml, anticoagulant: K_2_EDTA) were collected at 0, 0.083 (i.v. only), 0.25, 0.5, 1, 2, 4, 6, 8, 24, 48, and 72 hours post-dosing.

The subsequent pharmacokinetic and toxicology studies of **17** were conducted using solid dispersion and nanosuspension formulations in beagle dogs weighing 7 to 15 kg. The solid dispersion batch of compound **17** was dosed p.o. at 15 mg/kg of the active ingredient. The nanosuspension formulation was dosed in 1% hydroxypropyl methylcellulose, 0.2% sodium dodecyl sulfate. Blood samples were collected at 0, 0.25, 0.5, 1, 2, 4, 7, 24, 48, 72, 96, and 168 hours for the analysis of intact prodrug and metabolite **7** in plasma and triphosphate in PBMCs.

The toxicology studies were performed at Charles River Laboratories Preclinical Services Montreal (Sherbrooke, Canada) under the sponsor of Novartis. Male and female Wistar Han rats (Charles River, Raleigh, NC) and beagle dogs (Marshall BioResources, North Rose, NY) were used. The solid dispersion batch of **17** was prepared and administered daily by oral gavage (20 mg/kg for rats or 12 mg/kg for dogs) for up to two weeks at doses of 30 (dog only), 100, 300, and 1000 (rat only) mg/kg/day. Blood samples (anticoagulant: K_2_EDTA) were collected on day 1 and 14 (0.5, 1, 3, 7, and 24 hours post-1^st^-dose for dogs) for the analysis of intact prodrug and metabolite **7** in plasma and triphosphate **6** in PBMCs.

Prior to this study, a rising dose toxicology study had been conducted at single doses of 10, 30, 100 and 300 mg/kg. Assessment of the dose proportionalities of the plasma intact prodrug and metabolite **7** as well as the PBMC triphosphate were obtained for the toxicokinetic data of this rising dose study.

AG129 mice (lacking IFN-α/β and IFN-γ receptors (31)) was obtained from Biological Resource Center (BRC), Singapore. Male and female AG129 mice aged 8 to 14 weeks (weighing 20-30 grams) were used. Infection of DENV-2 (strain TSV01) was given intraperitoneally (500 µl, 1.4 × 10^7^ pfu/ml). The solid dispersion batch of **17** was dosed p.o. immediately after infection. The doses given were 10, 30, 100, and 300 mg/kg twice daily for 3 consecutive days. Blood samples (anticoagulant: K_2_EDTA) for pharmacokinetic analysis of the intact prodrug and nucleoside metabolite **7** in plasma were collected on day 1 and 3 post-infection (1, 3, 6, 24, 48, 50, 52, 55, and 72 hours post-1^st^-dose). Pooled blood samples of 6 mice were collected at the terminal time point for PBMC triphosphate analysis by LC-MS/MS. Plasma was also obtained from the terminal blood sample of each mouse for viremia read-out by plaque assay.

The non-compartmental pharmacokinetic parameters from various studies were calculated either using Watson LIMS (Thermo Fisher Scientific, Waltham, MA) or WinNonlin 5.01 (Pharsight Corporation, Mountain View, CA).

## RESULTS

### Chemistry and Structure-Activity/Property-Relationship (SAR/SPR)

We employed 2’-deoxy-2’-fluoro-2’-*C*-methylguanosine as a starting point and investigated a suitable prodrug moiety to effectively deliver the monophosphate into PBMCs for the treatment of dengue. From various types of nucleotide prodrugs, we focused our attention on 3’,5’-cyclic phosphoramidates (32). To access the cyclic phosphoramidate prodrugs, the nucleoside starting material, 6-*O*-alkyl-2’-deoxy-2’-fluoro-2’-*C*-methylguanosine **8** was prepared according to the literature (33). This guanosine analog **8** was reacted with pentafluorophenyl ester agent **9** in the presence of *tert*-BuMgCl as a base to afford the linear phosphoramidate product **10** as a mixture of diastereomers. The cyclization step was carried out by treatment of **10** with *t*-BuOK in DMSO to afford a diastereomeric mixture of cyclic products **11**, which were separated by RP-HPLC to give each single phosphorous stereoisomer (Scheme 1). The phosphorous stereochemistry of one of the cyclic phosphoramidates **17** was assigned as *R*p configuration as determined by single crystal X-ray analysis (Fig. 2). The X-ray structure of **17** indicates that the phosphoramidate moiety is *cis* oriented to the guanine base through H-bond formation between the carbonyl and 7-NH_2_. In the ^31^P NMR, the phosphorous peak of *R*p isomer **17** appeared at upper field than the corresponding *S*p isomer. This observation was applied to assign the phosphorous stereochemistry for the rest of the analogs.

**Scheme 1.**
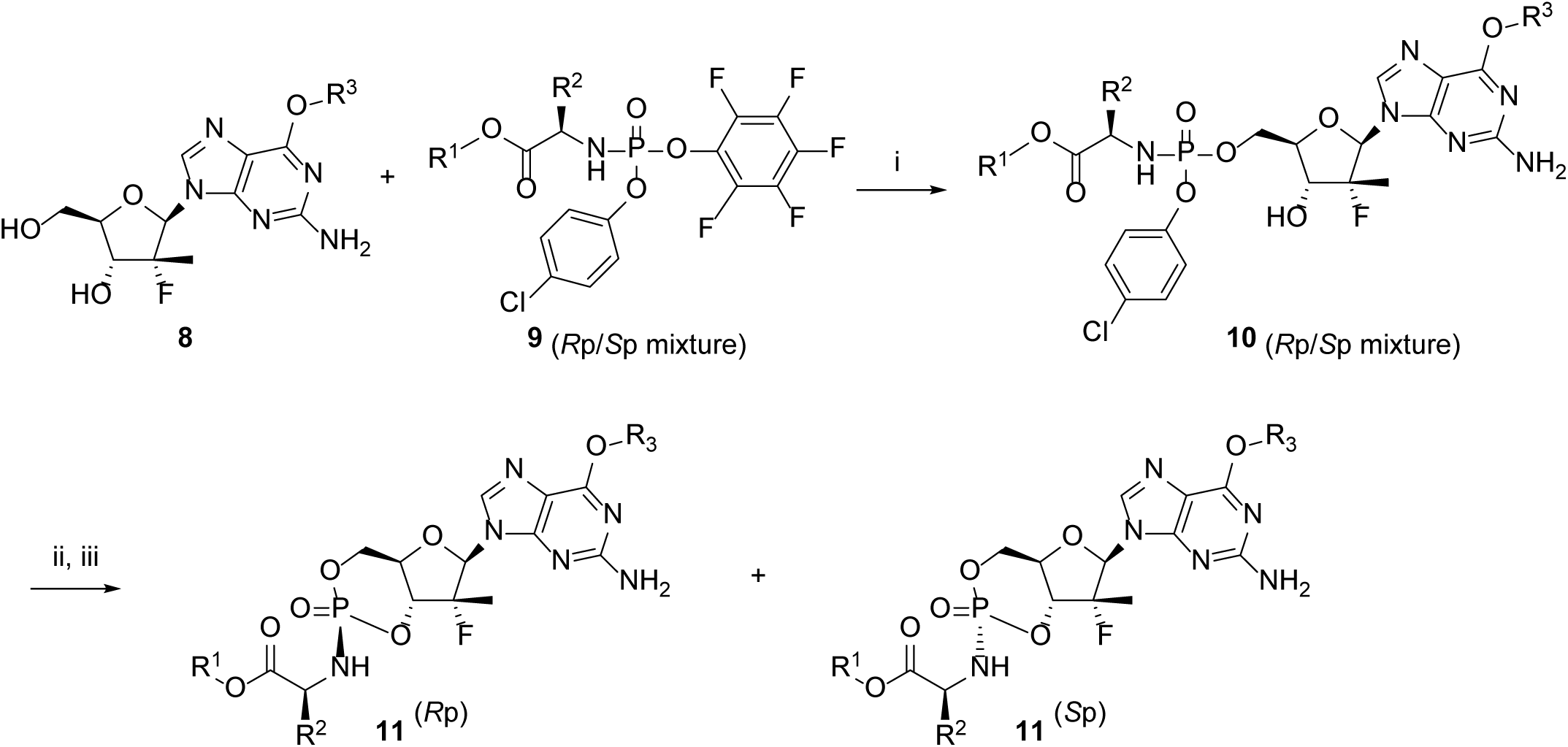
Synthesis of cyclic phosphoramidate prodrugs of 6-*O*-alkyl-2’-deoxy-2’-fluoro-2’-*C*-methylguanosine. Reactions conditions: (i) *t*-BuMgCl, THF; (ii) *t*-BuOK, DMSO; (iii) Separation by preparative reverse phase HPLC.

**FIG 2.**
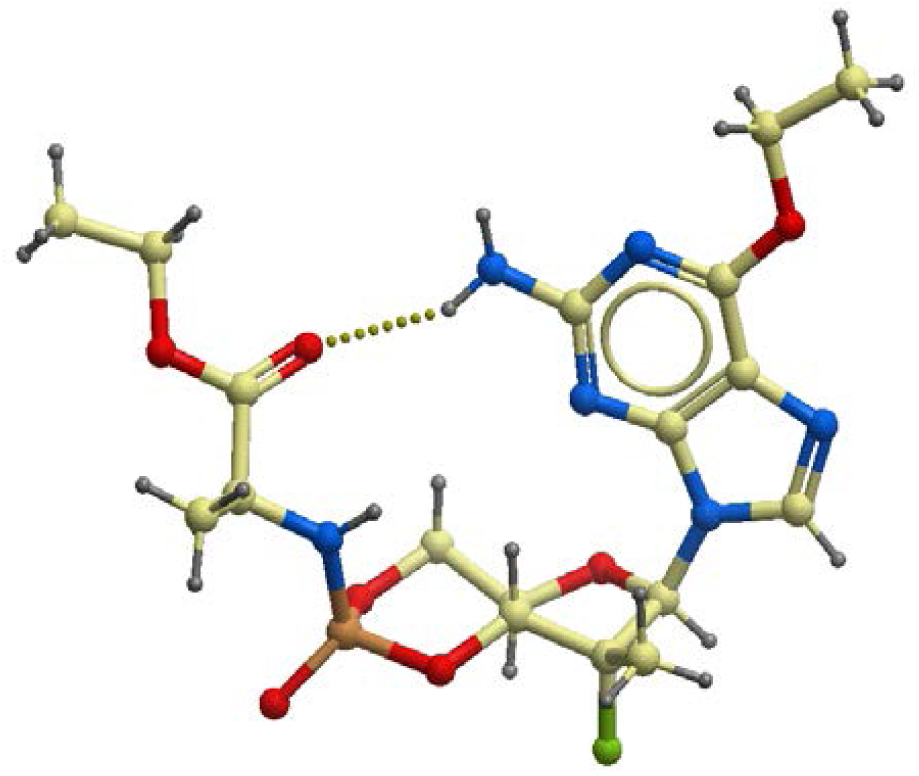
Single X-ray crystal structure of compound 17.

Over 150 cyclic phosphoramidates were synthesized with variations in the ester, amino acid, phosphorous stereoisomer, and *C*-6 substitutions on the guanine base. As highlighted in Table 1, the prodrugs were assessed by cellular anti-dengue activity, plasma stability, liver S9 stability, intestinal S9 stability to select the best prodrug with a balanced profile. Most of the prodrugs had over 120 min half-life in plasma and intestinal S9 fraction in higher species (dogs and human).

**TABLE 1.**
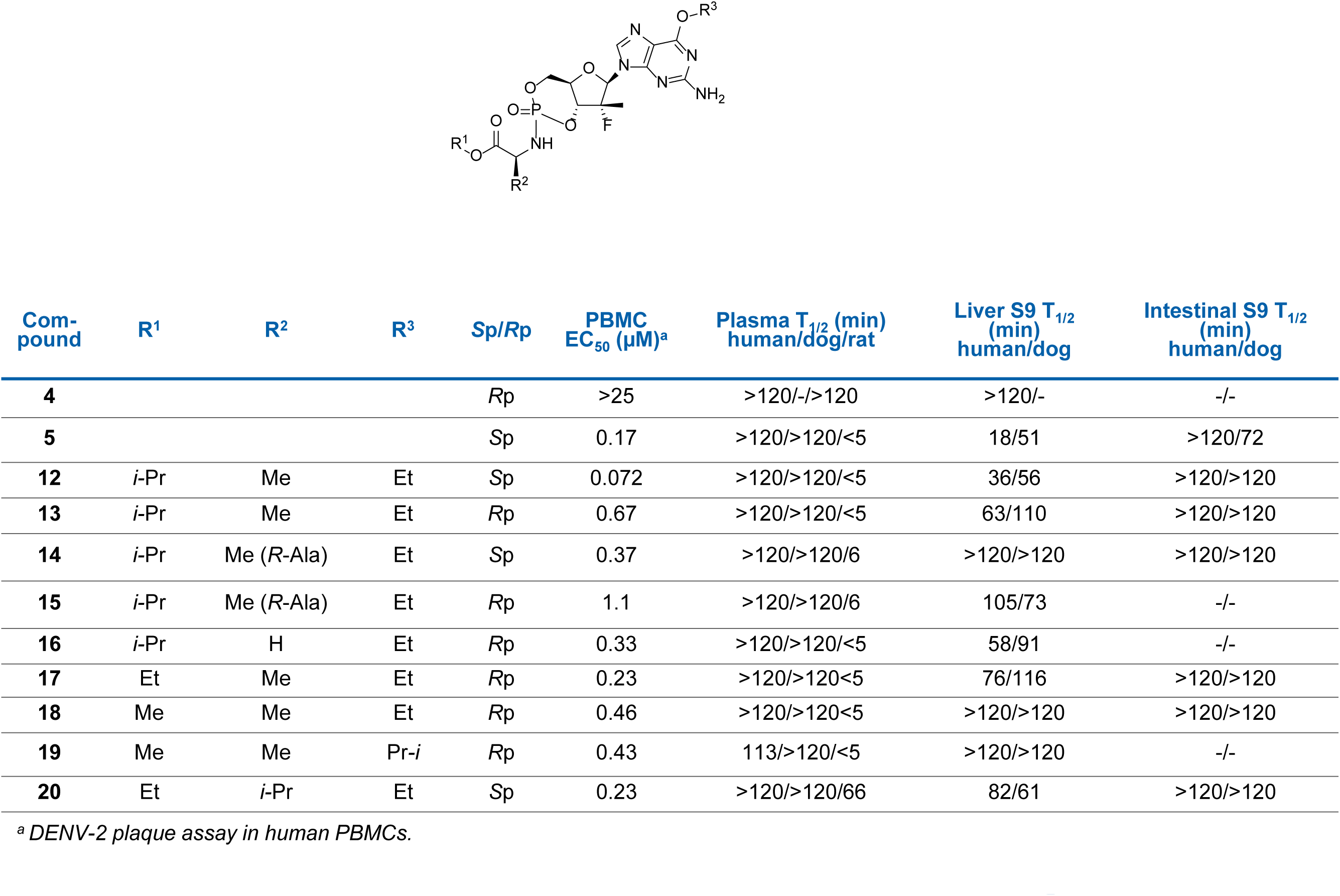
Anti-dengue activity and *in vitro* stability profile of cyclic phosphoramidate prodrugs of 6-*O*-alkyl-2’-deoxy-2’-fluoro-2’-*C*-methylguanosine.

The cyclic prodrug **12** had the same amino acid moiety (*S*-Ala-O*i*Pr) and phosphorus stereochemistry (*S*p) with **5** but showed 2-fold longer half-life than the linear prodrug **5** in human liver S9 fraction. The *R*p isomer **13** was much less potent with 2-fold longer liver S9 half-life as compared to the *S*p isomer **12**. The corresponding (*R*)-alanine analogs **14** and **15** led to reduced anti-dengue activity. The glycine analog **16** had similar potency and stability profile as the alanine prodrug **13**. The ethyl ester **17** showed 3-fold improvement in the potency with maintaining similar liver S9 stability. The methyl ester **18** had similar potency with good stability profile. The more lipophilic 6-*O*-*i*Pr analog **19** did not change anti-dengue activity and *in vitro* stability. The (*S*)-valine analog **20** was not as good as the (*S*)-alanine prodrug **12**. Based on the balance of antiviral activities and *in vitro* stability profiles, alanine-based prodrugs with different combination of esters and stereoisomers (**12, 14, 17, 18**) were selected for further characterization.

### Cyclic phosphoramidate prodrugs of 2’-deoxy-2’-fluoro-2’-*C*-methylguanosine convert to their corresponding triphosphate form in PBMCs in multiple species

Selected cyclic phosphoramidate prodrugs **12, 14, 17, 18** as well as the linear phosphoramidate PSI-353661 (**5**) were tested for its triphosphate concentration in PBMCs *in vitro*. A continuous incubation for 24 hours and prodrug concentration of 10 µM was chosen to allow the detection of the triphosphate forms at the linear phase (before saturation) of the enzymatic processes (see also Fig. 6).

**FIG 3.**
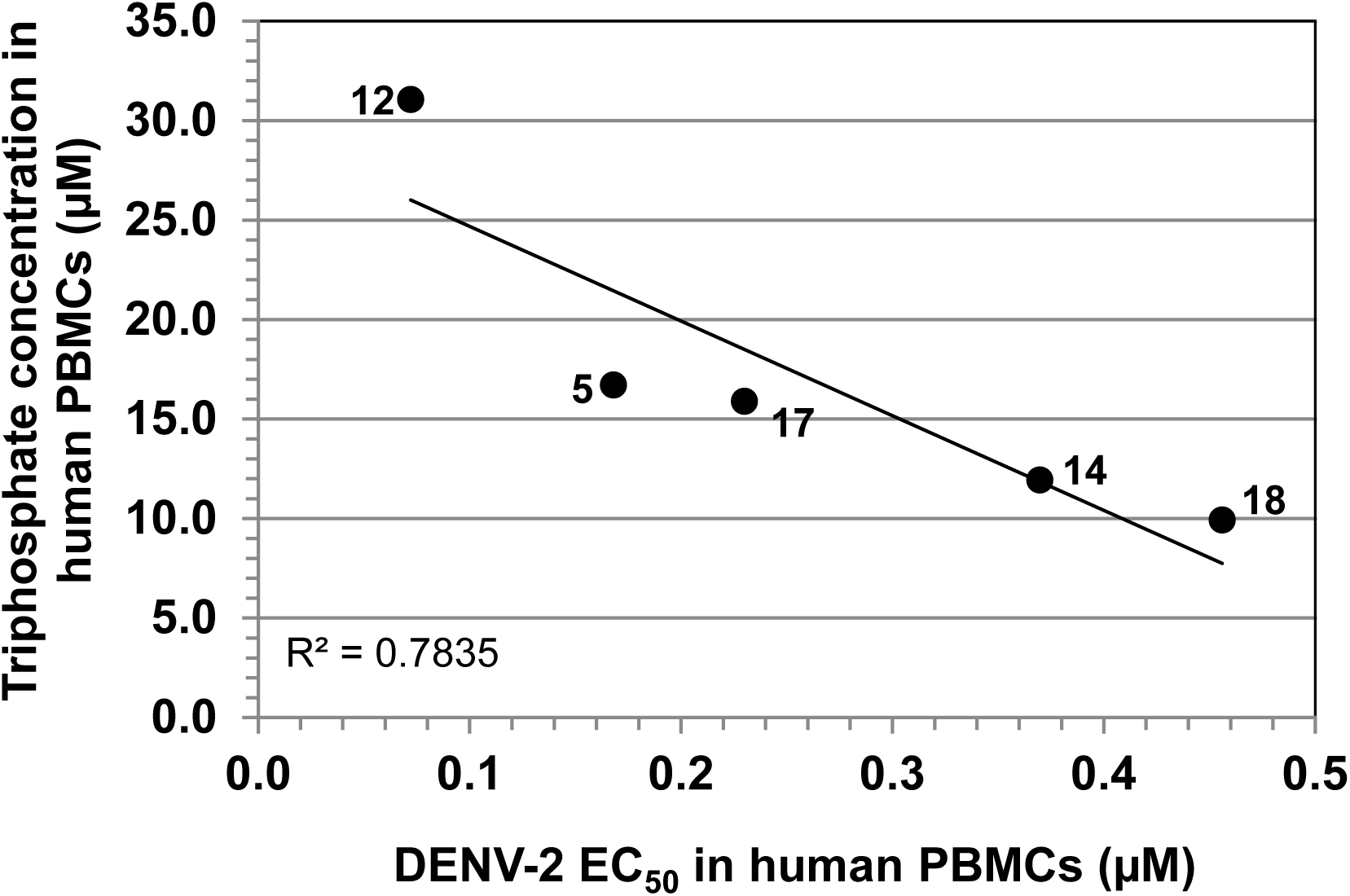
Correlation between triphosphate levels in human PBMCs *versus* potencies. Prodrugs were incubated in human PBMCs at 10 µM concentration. Data obtained were from at least two independent experiments.

**FIG 4.**
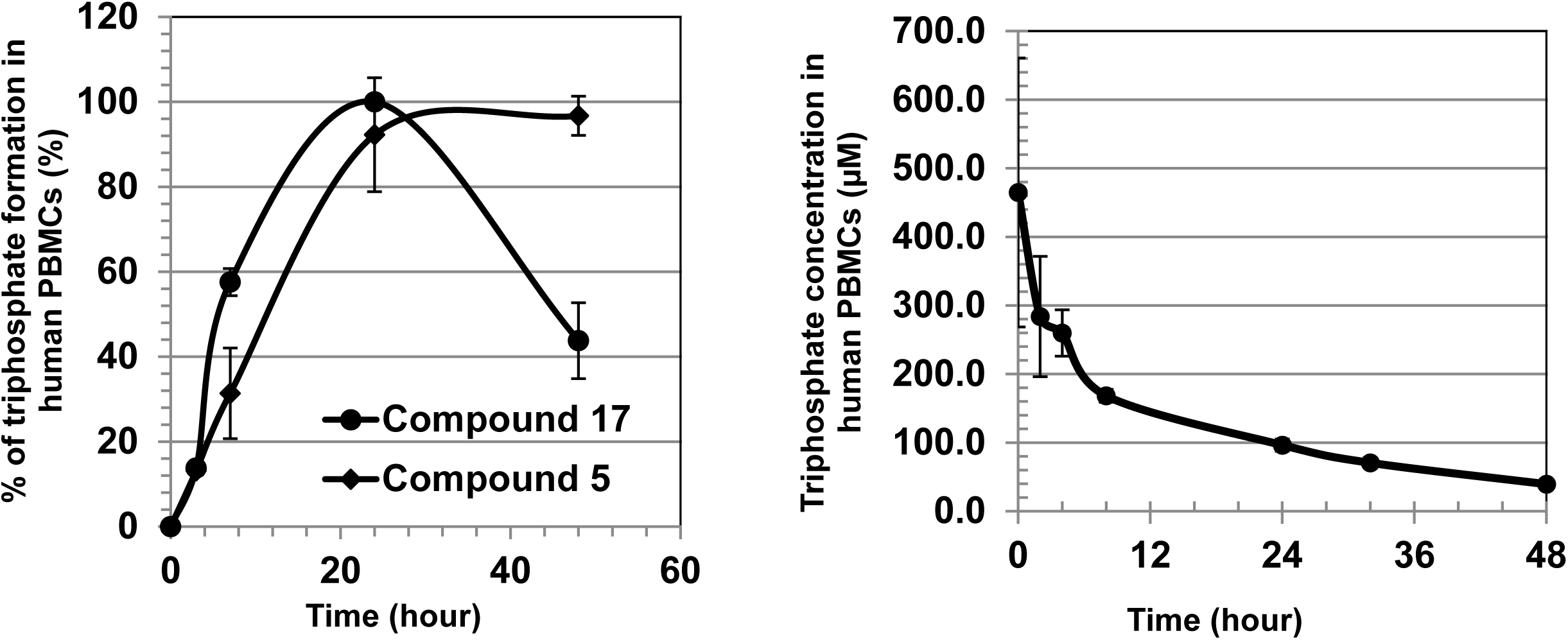
Triphosphate measurement of selected compounds in PMBCs **(A)** Triphosphate conversion kinetics of prodrug **17** and linear phosphoramidate analog **5**. The triphosphate was expressed as the percentage of the formation as compared to the highest triphosphate level achieved for each prodrug (n=3). **(B)** Triphosphate measurement of **6** with a sustained level (terminal t_1/2_ ∼20 hours) upon prodrug removal (n=3).

**FIG 5.**
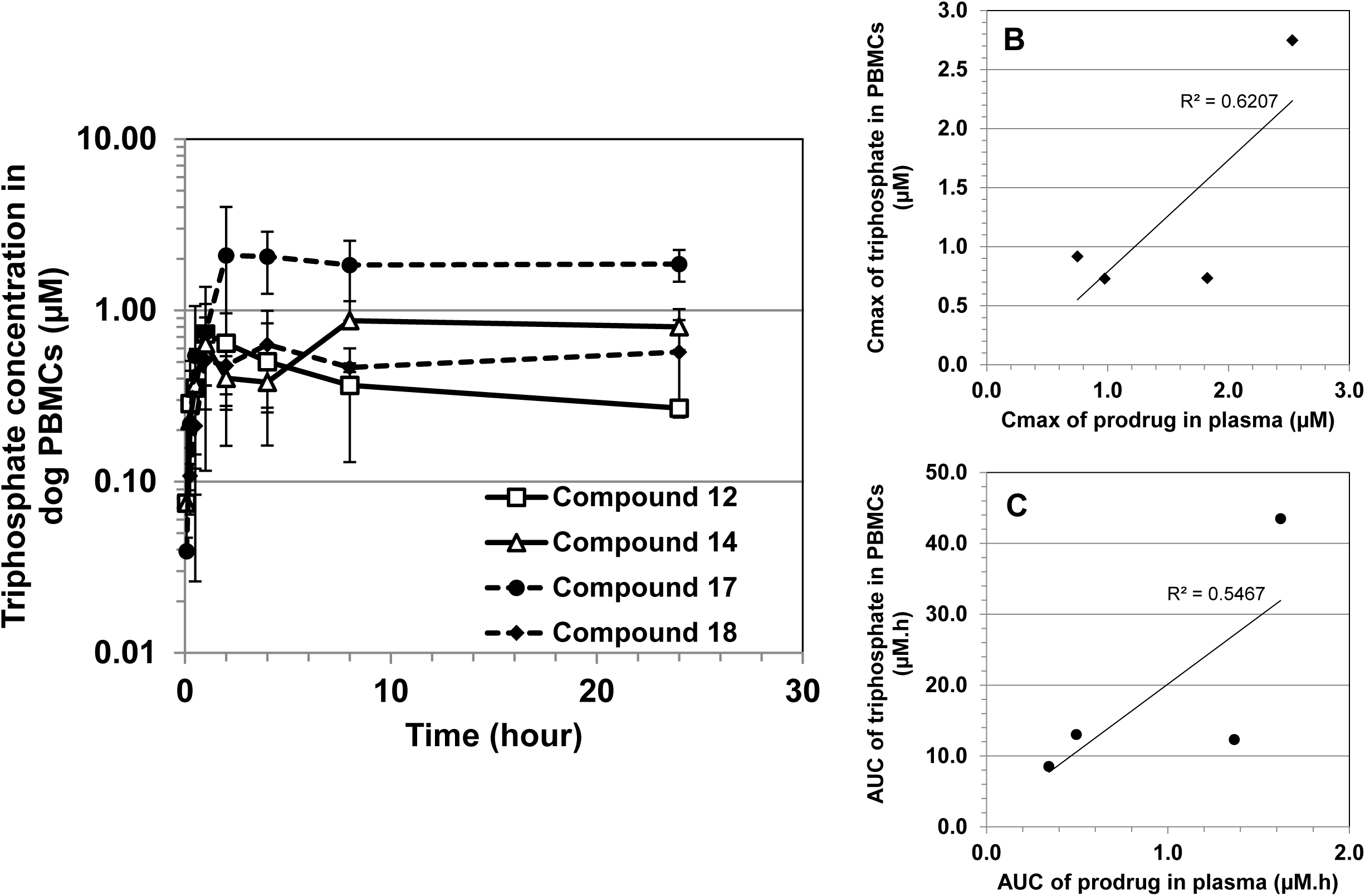
Selected prodrugs dosed orally (3 mg/kg) to beagle dogs (n=3). **(A)** Pharmacokinetic profiles of triphosphate metabolites in PBMCs. **(B)** Correlation between the C_max_ of prodrug in plasma and of triphosphate in PBMCs. **(C)** Correlation between the AUC of prodrug in plasma and of triphosphate in PBMCs.

**FIG 6.**
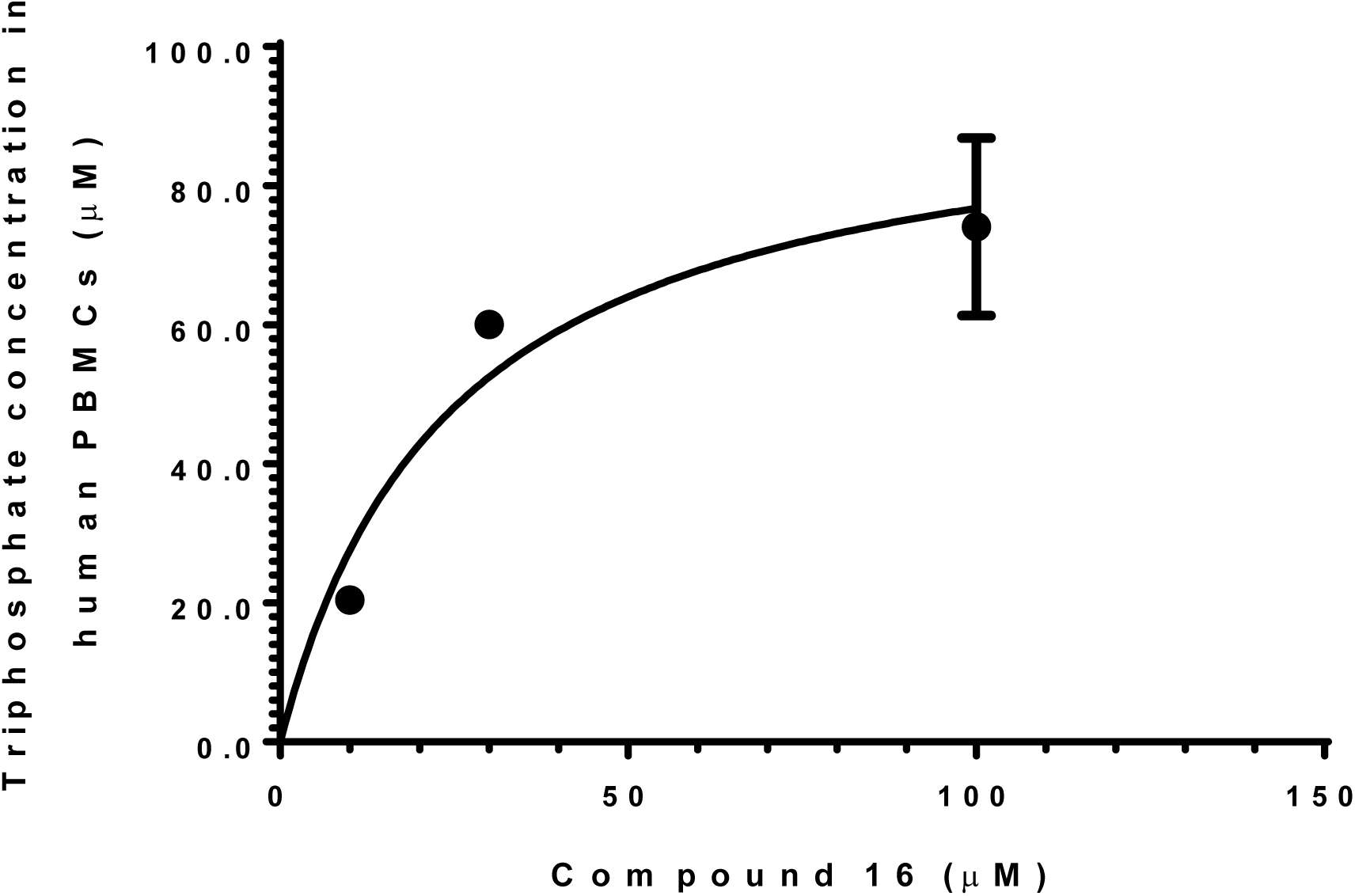
Incubation of compound **17** with increasing concentration in human PBMCs to determine TP_50_.

Different prodrugs converted to the same triphosphate at different rates and their potencies reflected the intracellular triphosphate concentrations reached. A trend of linear correlation was observed between triphosphate levels and potencies in human PBMCs (Fig. 3) and the most potent compound **12** showed the highest triphosphate level *in vitro*. Conversely, the intracellular triphosphate levels *in vivo* were determined not only by the triphosphate conversion in cells but also by the intact prodrug exposure in plasma. In this regard, compound **17** showed the highest *in vivo* triphosphate level in dog PBMCs instead of **12**. Compound **17** was then assessed for its triphosphate conversion in multiple species *in vitro*. Table 2 demonstrated that the prodrug was converted to different levels of triphosphate in the PBMCs of all the species relevant for safety and efficacy assessment (mouse, rat, dog, monkey, and human).

**TABLE 2.** Prodrug **17** were converted to triphosphate in the PBMCs from multiple species. Data were at least n=3.

| Compound 17,<br>incubation<br>concentration<br>($\mu\text{M}$) | Triphosphate concentration in PBMCs of multiple species ($\mu\text{M}$) | | | | |
| --- | --- | --- | --- | --- | --- |
|  | Mouse | Rat | Dog | Monkey | Human |
| 10 | $8.5 \pm 2.5$ | n.d. | $2.8 \pm 0.4$ | $39.2 \pm 14.5$ | $15.9 \pm 10.6$ |
| 100 | $53.4 \pm 8.4$ | $8.8 \pm 0.5$ | $13.8 \pm 1.8$ | $22.6 \pm 6.4$ | $59.7 \pm 19.4$ |

*n.d. = not determined*

### Cyclic phosphoramidate prodrug of 2’-deoxy-2’-fluoro-2’-*C*-methylguanosine shows similar triphosphate conversion kinetic as the linear phosphoramidate analog and sustained triphosphate level

The conversion of a phosphoramidate prodrug to its active triphosphate form is a multi-steps biotransformation process (34-37). To assess the kinetics of triphosphate formation and the impact of the cyclic phosphoramidate moiety, compound **17** (cyclic phoshoramidate) and **5** (linear phosphoramidate) were incubated in human PBMCs and the intracellular triphosphate concentration was measured at different time points. Both **17** and **5** showed similar kinetic in triphosphate formation (Fig. 4A). Furthermore, the intracellular triphosphate formed was sustained upon prodrug removal (Fig 4B, terminal half-life ∼20 hours).

### Cyclic phosphoramidate prodrugs of 2’-deoxy-2’-fluoro-2’-*C*-methylguanosine show different pharmacokinetic profiles and triphosphate levels in dogs

Compounds **12, 14, 17, 18** were dosed i.v. (0.5 mg/kg) and p.o. (3 mg/kg) in dogs to determine their pharmacokinetic parameters (Table 3). The triphosphate concentration in PBMCs was quantified and a trend of correlation between the C_max_ or AUC of the prodrug *versus* the triphosphate was observed (Fig. 5B-C and Table 3). Compound **17** showed the highest intracellular triphosphate level in PBMCs after oral dosing, thus it was selected for further characterization (Fig. 5A).

**TABLE 3.** Pharmacokinetic parameters of selected compounds dosed *i*.*v*. (0.5 mg/kg) and *p*.*o*. (3 mg/kg) to beagle dogs (n=3). Intact prodrug and the major metabolite, nucleoside **7**, were monitored in plasma, while the triphosphate metabolite **6** was measured in PBMCs from the *p*.*o*. study.

| Route |  | Unit | Intact prodrug |  |  |  | Nucleoside 7 (R=H) |  |  |  | Triphosphate 6 |  |  |  |
| --- | --- | --- | --- | --- | --- | --- | --- | --- | --- | --- | --- | --- | --- | --- |
| Prodrug |  |  | 12 | 14 | 17 | 18 | 12 | 14 | 17 | 18 | 12 | 14 | 17 | 18 |
| *i.v.* | **V<sub>dss</sub>** | L/kg | 0.4 |  | 0.5 | 0.5 |  |  |  |  |  |  |  |  |
|  | **ER\*** | % | >100 | --** | 67 | 45 |  |  |  |  |  |  |  |  |
|  | **T<sub>1/2</sub>** | h | 0.1 |  | 0.3 | 0.3 | 4.0 | 4.0 | 3.4 | 3.9 |  |  |  |  |
| *p.o.* | **C<sub>max</sub>** | μM | 1.0 | 0.8 | 2.5 | 1.8 | 0.8 | 0.8 | 0.5 | 1.3 | 0.7 | 0.9 | 2.7 | 0.7 |
|  | **T<sub>max</sub>** | h | 0.08 | 0.25 | 0.08 | 0.25 | 0.5 | 0.5 | 2 | 1 | 1.0 | 16 | 11 | 4.7 |
|  | **AUC<sub>inf</sub>** | μM.h | 0.3 | 0.5 | 1.6 | 1.4 | 4.4 | 4.4 | 4.6 | 7.1 | 8.5 | 13.0 | 43.5 | 12.3 |
|  | **F** | % | 25 | --* | 44 | 24 |  |  |  |  |  |  |  |  |

\*ER = hepatic extraction ratio, obtained from clearance data from the *i.v.* study. Dog liver blood flow of 42 mL/min/kg was used.
\*\**i.v.* study not performed

### Compound 17 shows pan-serotype and good antiviral activities in multiple cell lines

The triphosphate **6** was confirmed to be a potent inhibitor of dengue RdRp with IC_50_ of 1.1 µM in the Dengue 4 RdRp assay.

The activities of **17** against all the four DENV serotypes were examined in primary human PBMCs and in cell lines other than PBMCs. Compound **17** was active against all DENV serotypes as shown in Table 4. It also showed good activity in multiple cell lines which may be important for *in vivo* efficacy as DENV infection shows a broad tissue tropism (38).

**TABLE 4.** Activity of compound **17** across multiple serotypes and cell lines

| Assay type | Activity | Units |
| --- | --- | --- |
| PBMC DEN1 48h assay (EC<sub>50</sub>), | 0.18 ± 0.06 | μM |
| PBMC DEN2 48h assay (EC<sub>50</sub>) | 0.23 ± 0.04 | μM |
| PBMC DEN3 48h assay (EC<sub>50</sub>) | 0.36 ± 0.33 | μM |
| PBMC DEN4 48h assay (EC<sub>50</sub>) | 0.37 ± 0.14 | μM |
| THP-1 DEN2 assay (EC<sub>50</sub>) | 0.46 ± 0.20 | μM |
| KU812 DEN2 high content imaging assay (EC<sub>50</sub>) | 1.41 | μM |
| K562 DEN2 assay (EC<sub>50</sub>) | 2.79 ± 0.22 | μM |
| 293T DEN2 assay (EC<sub>50</sub>) | 3.40 | μM |
| Huh-7 DEN2 replicon, luciferase assay (EC<sub>50</sub>) | 1.73 ± 1.06 | μM |

### TP_50_ -determination of exposure target for efficacy

An exposure target (TP_50_) was established to assess the efficacy of **17** *in vivo*. TP_50_, the intracellular triphosphate concentration at which 50% of the virus replication is inhibited, was determined in human PBMCs. The TP_50_ value of compound **17** obtained from 3 independent experiments (triplicate measurement in each experiment) is 0.78 ± 0.43 µM. Fig. 6 shows that the prodrug incubation followed Michaelis-Menten kinetic: saturation of the triphosphate conversion process was observed.

### Proof-of-concept in mouse model

Compound **17** (given p.o. at 10, 30, 100, and 300 mg/kg twice daily for 3 days) was assessed in the AG129 viremia mouse model (31). The mice infected with DENV-2 (strain TSV01) leads to viremia which peaks on day 3 post-infection.

The intact prodrug was not detectable in the plasma samples due to high esterase activities in rodents. Nucleoside metabolite **7**, the final biotransformation product in plasma, was detected. Table 5 shows the pharmacokinetic parameters obtained and a trend of dose-proportionality. The PBMC triphosphate terminal concentration was quantified from a pool of 6 mice. On day 3, the intracellular triphosphate levels of the 30 and 100 mg/kg twice daily groups were 0.39 and 1.43 µM, respectively. The terminal triphosphate concentration reached TP_50_ (0.78 µM) for the 100 mg/kg but not for the 30 mg/kg group and showed efficacy at 100 and 300 mg/kg twice daily as it reduced viremia significantly by 28- and 54-fold (or 1.6- and 2.2 log), respectively. The compound was not efficacious at 10 and 30 mg/kg twice daily (viremia reduction was 3- and 4-fold, respectively, and not significant) (Fig. 7).

**TABLE 5.** Pharmacokinetic parameters (n = 3 animals) upon twice-daily oral dosing of compound **17** in the infected mouse model. The intact prodrug level was not determined. Major metabolite nucleoside **7** was monitored at 1, 3, 6, and 24 hours post 1^st^ and 5^th^ dose in plasma. Terminal concentration of triphosphate metabolite **6** was measured in pooled PBMCs from the 30 and 100 mg/kg groups.

|  | Unit | Day | 10 mg/kg | 30 mg/kg |  | 100 mg/kg |  | 300 mg/kg |
| --- | --- | --- | --- | --- | --- | --- | --- | --- |
| Metabolite |  |  | 7 | 7 | 6 | 7 | 6 | 7 |
| C<sub>72</sub> | μM | 3 | 0.02 | 0.1 | 0.39 | 0.2 | 1.43 | 1.9 |
| C<sub>max</sub> | μM | 1 | 0.3 | 0.9 | -- | 2.5 | -- | 1.9 |
|  |  | 3 | 0.3 | 0.8 |  | 1.4 |  | 2.4 |
| AUC<sub>inf</sub><sup>*</sup> | μM.h | 1 | 2.7 | 7.7 |  | 23.2 |  | 24.4 |
|  |  | 3 | 3.3 | 9.5 |  | 20.6 |  | 29.6 |

*\*AUC<sub>(0-24)</sub> was used for 300 mg/kg group as extrapolation to infinity was over-predicted*

**FIG 7.**
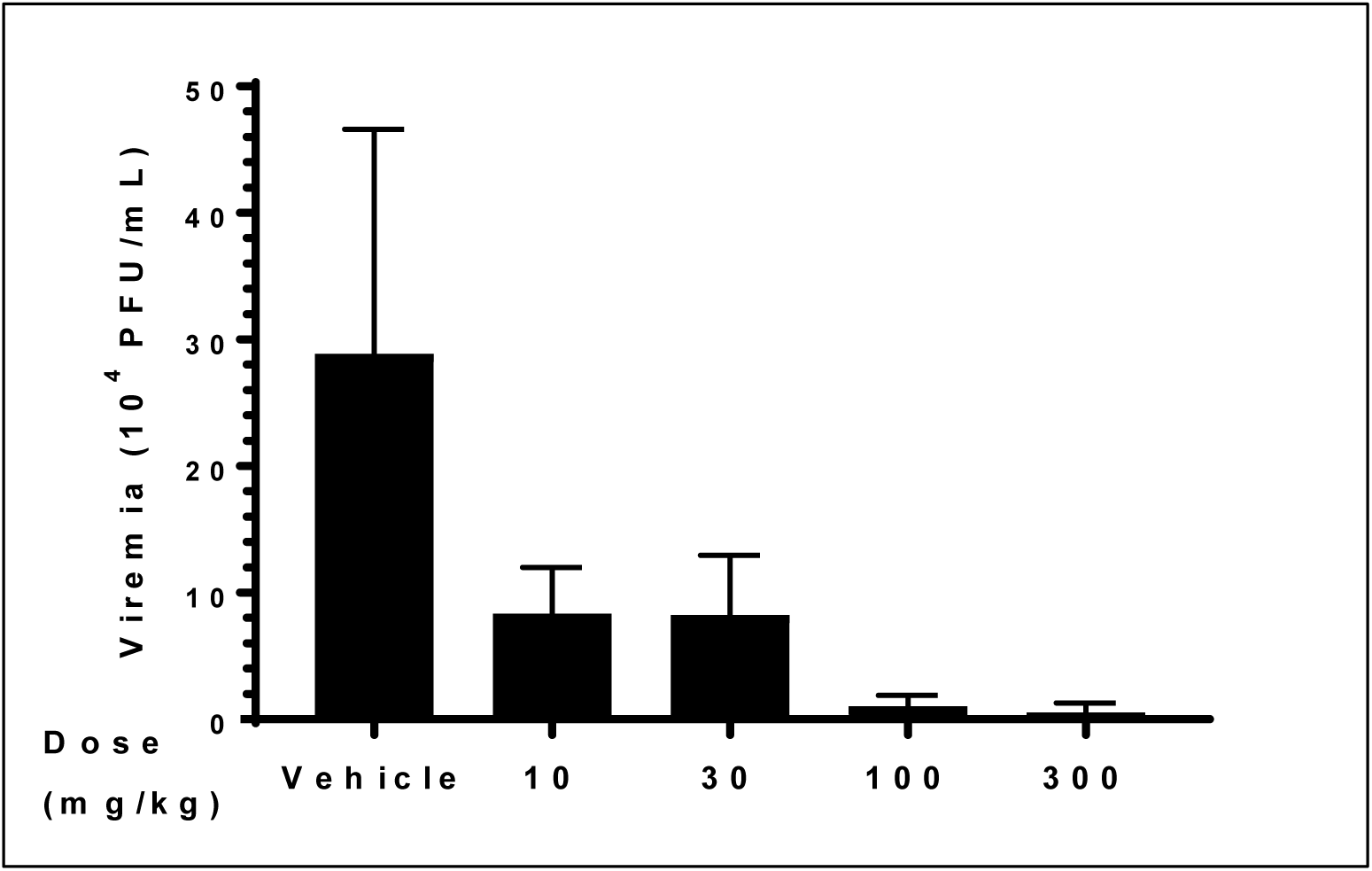
Efficacy study in viremia mouse model. Each group contains 6 mice. Compound **17** reduced viremia by 3-, 4-, 28- and 54-folds at 10, 30, 100 and 300 mg/kg twice daily, respectively. The viremia reduction at 100 and 300 mg/kg twice daily are significant (p<0.0001). The difference in the viremia reduction between 30 and 100 or 300 mg/kg twice daily are also significant (p<0.01 or p<0.0001, respectively).

### Formulation work and dose escalation study

Solution, nanosuspension, and solid dispersion formulations were tested by dosing compound **17** orally at 15 mg/kg to beagle dogs (Table 6). Solid dispersion showed the highest oral bioavailability.

**TABLE 6.** Pharmacokinetic parameters of compound **17** dosed p.o. at 15 mg/kg in different formulation to beagle dogs. Solid dispersion formulation has the highest oral bioavailability.

|  | Unit | Solution | Nanosuspension | Solid dispersion |
| --- | --- | --- | --- | --- |
| C<sub>max</sub> | μM | 5.5 | 0.9 | 16.4 |
| T<sub>max</sub> | h | 0.58 | 0.8 | 0.25 |
| AUC<sub>inf</sub> | μM.h | 4.4 | 1.5 | 12.2 |
| F | % | 24 | 8.5 | 68 |

Using the optimized formulation, the dose-proportionality was assessed in dogs (10, 30, 100, 300 mg/kg). A dose-proportionality was observed for the intact prodrug and metabolite **7** in plasma (Fig. 8A-B and Table 7). The intact prodrug had a short half-life (<1 hour), while the metabolite **7** has a long half-life (9-13 hour) and constituted the major metabolite in dogs. Other phosphoramidate intermediates were also detected in plasma but at lower levels and much shorter half-lives (data not shown).

**TABLE 7.**
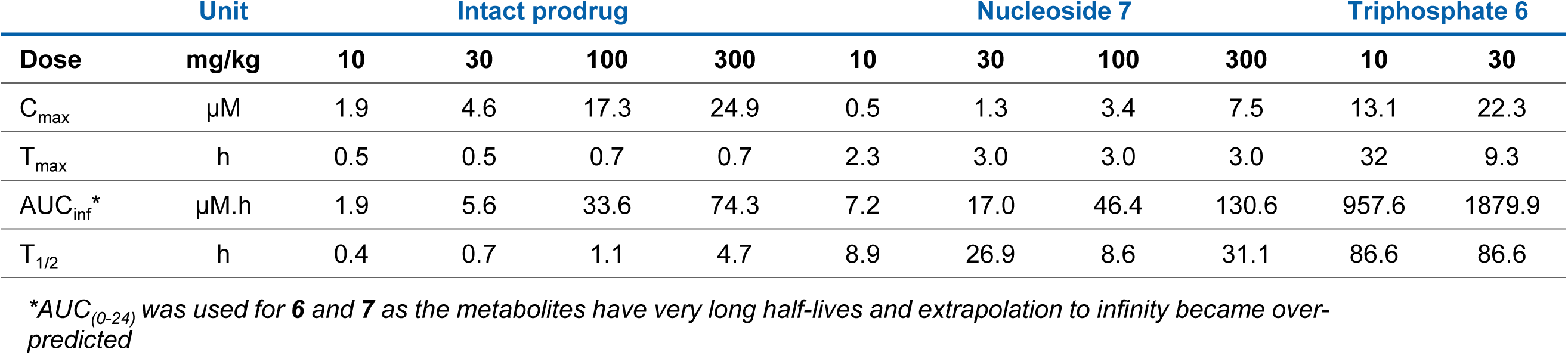
Pharmacokinetic parameters of compound **17** dosed p.o. at 10, 30, 100, and 300 mg/kg in solid dispersion formulation to beagle dogs. A trend of dose-proportionality observed.

**FIG 8.**
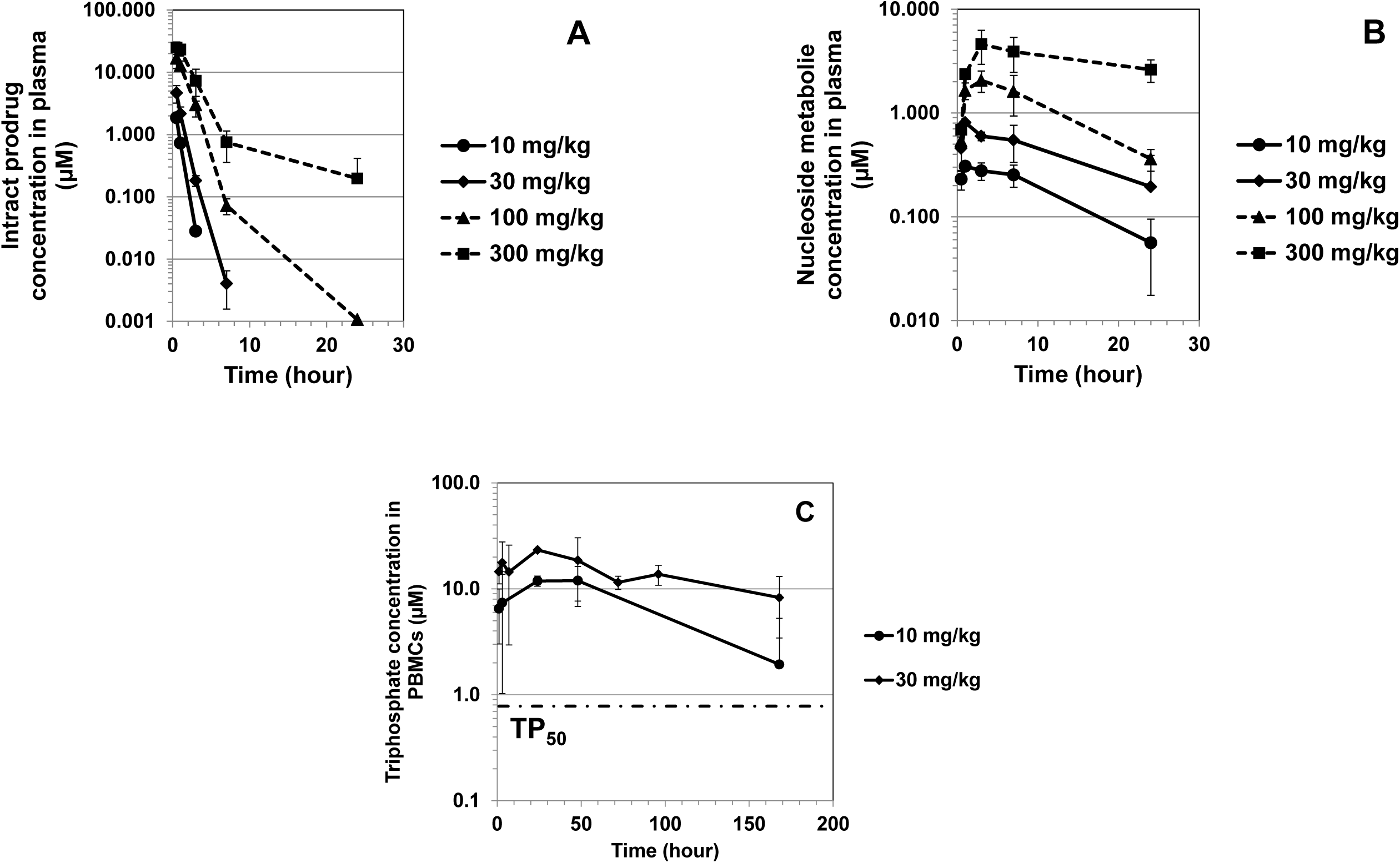
Pharmacokinetic profiles of the intact prodrug (**A**) major metabolite nucleoside **7** in plasma (**B**) and triphosphate metabolite **6** in PBMCs (**C**) upon raising doses of compound **17** in beagle dogs (n=3). The triphosphate concentration in PBMCs was determined only from the 10 and 30 mg/kg groups. At 10 mg/kg, the triphosphate level in PBMCs has exceeded TP_50_.

Upon single oral dose of **17**, a prolonged exposure of triphosphate **6** in PBMCs was observed. The triphosphate half-life was 3.5 days in dogs. A trend of dose-proportionality was observed albeit the high variability of the triphosphate levels (Fig. 8C and Table 7).

### *In vitro* and *in vivo* safety assessment

Compound **17** was assessed in multiple cell lines (HepG2, THP-1, MT-4 (27) and PC-3 (39)) as well as in various *in vitro* biochemical assays including the mini-Ames test for genotoxicity, hERG (human ether-a-go-go related gene) channel for cardiovascular toxicity, CYP450 inhibition for drug-drug interaction, micronucleus assay for mutagenicity, and various receptors, ion channels, and kinase profiles. The compound did not show significant inhibition in any of these assays. PSI-353661 (**5**) has been assessed in various cytotoxicity assays including in HepG2, huh-7, and BxPC3 and the compound shows CC_50_ ≥80 µM (35). We demonstrated here that a modification of the prodrug moiety (from linear to cyclic phosphoramidate) does not change the *in vitro* toxicity profiles.

With the clean *in vitro* profile, compound **17** was administered to rats (30, 100, 300, and 1,000 mg/kg/day) and dogs (30, 100, and 300 mg/kg/day) for up to 14 days for *in vivo* toxicology evaluation. The compound was tolerated in rats when given up to 1,000 mg/kg/day for 14 consecutive days. Unfortunately, compound **17** was poorly tolerated in dogs. On day 7-9, significant findings in the lung (inflammation and hemorrhage) led to severe decline in the canine health, moribund and necessary termination of some animals from the top dose group. The findings were dose-related and mild pulmonary inflammation and hemorrhage were already observed in one of the six dogs in the 30 mg/kg/day group and no observed adverse effect level (NOAEL) was not achieved in the dogs. Table 8 shows the triphosphate levels in dog PBMCs obtained on day 1 and day 14 (only from the 30 mg/kg/day group). A trend of dose-proportionality was observed between 30 and 100 mg/kg/day, but the proportionality diminished between 100 and 300 mg/kg/day (Table 7). No accumulation was observed between day 1 and day 14 triphosphate levels at 30 mg/kg/day.

**TABLE 8.**
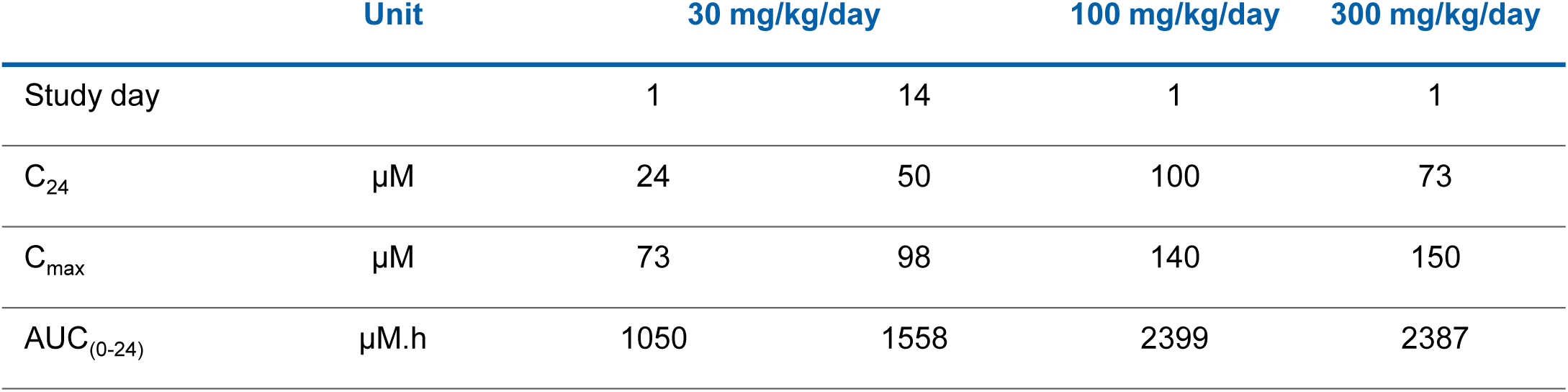
Triphosphate concentration in PBMCs upon multiple oral dosing of compound **17** in beagle dogs (n = 3 females and 3 males).

## DISCUSSION

Our objective is to develop an oral dengue drug that is efficacious and safe. Nucleoside analogs offer several advantages as dengue drug candidates as they target an essential viral specific enzyme RdRp (5), have pan-serotypic activity and high resistance barrier (8). As PBMCs is one of the major viral replication sites (24), the active triphosphate concentration in PBMCs was used as a pharmacological marker for efficacy in addition to EC_50_ value generated in PBMCs. The same measurement of triphosphate concentration in the target organs has also been used as a pharmacological marker for HIV (40).

Several successful examples of nucleoside antivirals have been developed for HIV, HSV, HBV, and HCV therapeutic areas (6). In dengue, two different nucleosides have been reported i.e. NITD-008 and balapiravir. NITD-008 demonstrated good oral efficacy in the dengue model, but it was unable to progress to human clinical trials due to insufficient safety profile (10). Balapiravir was repurposed from HCV in a phase II dengue clinical trial, but failed to reduce viral load in the patients (13). Although the reasons remain to be fully understood, one potential reason could be the state of PBMC dengue viral infection (12).

In an attempt to bypass the rate-limiting phosphorylation step, we explored the potential of monophosphate prodrug approach, starting from guanosine-based nucleoside analogs. Monophosphate prodrugs have been developed to deliver nucleoside monophosphates directly into the cells, resulting in higher intracellular level of the active triphosphates (16, 18). Examples of monophosphate prodrug strategies for 2′-deoxy-2′-fluoro-2′-*C*-methylguanosine were demonstrated by Pharmasset and exemplified by PSI-352938 (**4**) (21, 22) and PSI-353661 (**5**) (23) for HCV treatment. PSI-352938 passed preclinical safety assessment and was well-tolerated at doses of up to 1600 mg once daily in phase I study (18). In a later phase II study, compound **4** caused liver function abnormalities after 13 weeks dosing (41). Although PSI-352938 caused hepatotoxicity after chronic dosing, we reasoned that this could still be a good starting point for dengue since the toxicity is reversible and DENV treatment duration need only be a week or less (9). In addition, we suspected the triphosphate metabolite could be the reason for liver toxicity and hypothesized that modification of prodrug moiety could change the compound distribution and thus reduce liver toxicity. Our prodrugs are specifically designed to maximize the stability in GI, liver, and systemic circulation before penetrating into PBMCs. Once inside, the intracellular enzymes in PBMC would unmask the prodrug moieties easily, allowing further metabolism to the active triphosphate. We used *in vitro* stability in liver and intestinal S9 fractions, plasma stability together with cellular anti-dengue activity in PBMCs to find the most balanced compound. Indeed when tested *in vivo*, compound **17** generated the highest level of triphosphate in dog PBMCs. Indeed, the major organ for toxicity has shifted from liver to lung, possibly due to much higher systemic and lung exposure.

The common guanosine-based nucleoside triphosphate **6** of PSI-352938 and PSI-353661 showed potent inhibition for dengue RdRp with IC_50_ value of 1.1 µM. PSI-352938 (**4**) and PSI-353661 (**5**) also displayed inhibition of DENV in Huh-7 replicon cells with EC_50_ values of 0.76 µM and 0.044 µM respectively. The corresponding free nucleoside, 2’-deoxy-2’-fluoro-2’-*C*-methylguanosine **7** was inactive (EC_50_ >50 μM), suggesting the addition of first phosphate addition is the rate limiting step in Huh7 cells. On the other hand, PSI-352938 (**4**) was not active in the dengue human PBMC plaque assay since the first step of activation of **4** was triggered by CYP3A4 mediated P-*O*-dealkylation (37), which is absent in PBMCs. Finally, PSI-353661 (**5**) exhibited potency against dengue with EC_50_ of 0.17 µM in PBMCs. Compound **5** was not ideal for dengue as it had a short half-life in liver S9 fraction (<20 min), and was designed for targeting the liver and not for systemic distribution as required. We therefore pursued a suitable prodrug moiety to balance the stability profiles, particularly in liver S9 and EC_50_ in PBMCs.

From various types of nucleotide prodrug, we became interested in 3’,5’-cyclic phosphoramidates (32). This particular prodrug could offer additional advantages as compared to the linear phosphoramidates like PSI-353661 (**5**) as they masked the 3’-free OH and reduced a degree of rotational freedom, potentially allowing for improved cell entry and prolonged metabolic stability in the liver. In addition, 3’,5’-cyclization eliminated the release of toxic aromatic alcohols like phenol and naphtol. Over 150 cyclic phosphoramidates were synthesized with variation in the ester, amino acid, phosphorous stereoisomer, and *C*-6 substitution on the guanine base. Based on the balance between antiviral activity and *in vitro* stability profiles, we selected alanine prodrugs with different combination of esters and stereoisomers (**12, 14, 17, 18**) for further characterization. We directly monitored the concentration of the active triphosphate **6** in PBMCs and observed a trend of linear correlation between potencies and PBMC triphosphate levels. This correlation is expected as the triphosphate is the active form inhibiting viral replication through termination of RNA-chain synthesis (7, 10).

*In vivo* pharmacokinetic profiling for the selected prodrugs **12, 14, 17, 18** were performed in dogs using i.v. at 0.5 mg/kg and p.o. at 3 mg/kg to establish plasma clearance, oral absorption and triphosphate loading in PBMCs over 24 hours. Prodrug **12** had the least stability (the highest clearance) both *in vitr*o in liver S9 and *in vivo*. All the prodrugs had good solubility (>0.4 g/L from high-throughput equilibrium solubility) and low-to-moderate permeability (data not shown) and the *in vivo* pharmacokinetic profiles obtained were similar. Overall, a good *in vitro – in vivo* correlation (IVIVC) was observed for all the compounds. Compound **17** showed the highest level of triphosphate in PBMCs (about 3-fold), albeit with high inter-animal variabilities, confirming our hypothesis that a balanced S9 and plasma stability combined with potent antiviral activity led to the highest level of triphosphate in PBMC. The fact that multiple host enzymes were needed to convert the prodrug to pharmaceutically active triphosphate may explain the higher variability among animals tested. It was also known that small modification of the phosphoramidate prodrugs could change the triphosphate conversion dramatically (42). Due to the highest loading of the triphosphate in PBMCs when dosed *in vivo*, **17** was selected for further characterization.

To assess the relevance of various animal models for pharmacokinetic, efficacy, and safety evaluation of a nucleoside prodrug, we first tested compound **17** for triphosphate conversion in PBMCs of multiple species. In fact, species difference in triphosphate conversion has been well documented (36, 37, 42, 43). We demonstrated that **17** was converted to the active triphosphate in the PBMCs of all the relevant species, including mice and monkeys which enable compound assessment in the two different dengue animal models if needed (31, 44).

We also evaluated the behavior of triphosphate **6** in PBMCs in terms of conversion kinetic and half-life. Our data demonstrated that both cyclic and linear phosphoramidate prodrugs showed similar triphosphate conversion kinetics *in vitro*. Furthermore, the triphosphate level of **17** is sustained upon prodrug removal, with the half-life (∼20 hours) similar to other reported nucleoside triphosphates in lymphocyte or monocyte-derived cells (45, 46).

To define the minimum efficacious dose of prodrug **17**, we defined TP_50_ as the intracellular triphosphate concentration at which 50% of the viral replication is inhibited. TP_50_ is defined as the intracellular concentration of triphosphate yielded upon incubation of prodrug at its EC_50_ in human PBMCs. We found that the TP_50_ value of **17** (0.78 µM) was close to its IC_50_ (1.1 µM) obtained from the polymerase enzyme assay. Next, we wanted to assess if the level of triphosphate can be translated to efficacy *in vivo* using a dengue viremia mouse model (31). Due to the abundant carboxylesterases in the plasma of rodent species (36, 47), 100 and 300 mg/kg twice daily oral dosing for 3 days was needed to achieve at least 1 log viremia reduction (28- and 54-fold viremia reduction, respectively), while the efficacy was not observed at 10 and 30 mg/kg (only 3- and 4-fold viremia reduction, respectively). Pooled terminal blood samples were taken on day 3 post-infection from the 30 and 100 mg/kg groups and measured for the triphosphate concentration in PBMCs. The intracellular triphosphate concentration on day 3 reached TP_50_ (0.78 µM) for the 100 mg/kg group (1.43 µM), but not for the 30 mg/kg group (0.39 µM). Based on these *in vitro* and *in vivo* results, we defined here for the first time the minimum efficacious dose for a nucleos(t)ide prodrug as the dose that is required to maintain triphosphate concentration in PBMCs above TP_50_. By applying this principle to dog species, whose plasma stability is close to human, compound **17** reached TP_50_ already at 10 mg/kg (Fig. 8C).

Having been able to estimate minimum efficacious dose in dogs, the compound was prepared for preclinical toxicological evaluation *in vivo*. Upon physical form screening, **17** showed more than one crystal structures (polymorphism). A single crystal form was selected for its superior physico-chemical properties. Assessment of this form in dogs showed poor oral bioavailability (1%) from conventional suspension formulation (0.5% Tween80 and 0.5% methyl cellulose in water - data not shown). To enable high dose toxicology studies, a formulation work was conducted.

A high concentration solution formulation of compound **17** (≥10 mg/ml) with low total organic content (<30%) could not be achieved. The selected crystal form of **17** had low to medium aqueous solubility (<0.1 mg/ml in pH 3 to 6.8 and in bio-relevant media), low intrinsic dissolution rate and a low logP. A nanosuspension formulation was developed but resulted in low oral bioavailability (8.5%). Eventually, a solid dispersion formulation was feasible for high dose *in vivo* studies. Upon evaluation in dogs, the solid dispersion formulation improved the oral bioavailability (68%) from suspension (1%) as well as from the solution formulation (oral bioavailability 24% at similar dose) used in our early prodrug selection. Using this optimized formulation, compound **17** was assessed for dose-proportionality in dogs. The compound showed a trend of dose-proportionality for the intact prodrug and free nucleoside metabolite in plasma as well as for the triphosphate in PBMCs. The levels of triphosphate was sustained at a level above TP_50_ for at least one-week after a single dose of 10 mg/kg (half-life 3.5 days), making the compound potential for a single-dose cure.

Compound **17** was further assessed for safety in rat and dog toxicology studies. Although the stability of **17** in rat plasma is <5 min, the triphosphate could still be detected in PBMCs upon multiple oral dosing. An oral administration of **17** was tolerated up to 1,000 mg/kg/day when given for 14 consecutive days to wistar rats. A quantitative whole-body autoradiography (QWBA) study shows an extensive distribution to most rat tissues, except the central nervous system and testis, after a single-dose of 100 mg/kg [^14^C]compound **17** (data not shown). This observation is in agreement with our hypothesis that the change in prodrug moiety could lead to systemic distribution.

Next, compound **17** was administered orally to beagle dogs for up to two weeks at 30, 100, and 300 mg/kg/day. The compound was not tolerated at 100 and 300 mg/kg/day and clinical signs accompanied by weight loss were first observed in dogs in day 7; which was accompanied by early termination for these two groups. Liver was not the target organ for this compound, unlike the liver-targeting PSI-352938, as our prodrug moiety has changed the compound distribution. However, tubular degeneration of the kidneys, lung inflammation and hemorrhage were observed, among other findings.

Dose proportionality of the intact prodrug and nucleoside metabolite in plasma as well as the triphosphate in PBMCs was observed from 30 to 100 mg/kg/day, but was under proportional from 100 to 300 mg/kg/day (Table 7). The triphosphate reached very high exposure in PBMCs and there was no triphosphate accumulation observed upon multiple dosing of 30 mg/kg/day for 14 consecutive days. The triphosphate levels achieved in dogs were generally higher than the ones in rats at similar doses i.e. terminal concentration in PBMCs (day 15) was 10 µM in rats (data not shown) and 50 µM in dogs, which may be the reason for the clean finding in rats.

The pathology findings in dogs were dose-related and mild pulmonary inflammation and hemorrhage was already observed in one of the six dogs given 30 mg/kg/day of compound **17** for 14 consecutive days. Therefore, a no observed adverse effect level (NOAEL) was not achieved in this study. Due to the severity of the adverse findings and only partial reversibility of such findings at high doses, further development of **17** was not pursued.

Taken together, we have shown the potential of monophosphate prodrugs for dengue and demonstrated a suitable prodrug moiety to effectively deliver the monophosphate into PBMCs upon oral dosing. We have addressed the efficacy of monophosphate prodrugs by demonstrating a proof-of-concept in a mouse model. We establish TP_50_, the intracellular triphosphate concentration at which 50% of the virus replication is inhibited, as an exposure target and define the minimum efficacious dose as the one that is required to maintain triphosphate concentration in PBMCs above TP_50_. This concept could be universally applied and will be useful to evaluate the efficacy of any dengue nucleos(t)ide monophosphate prodrug.

## ACKNOWLEDGEMENT

The authors wish to thank Kwan Leung, David Beer, Paul Smith, Lv Liao, Margaret Weaver and Thierry Diagana for their helpful discussions to this project. Peter Wipfli, Ying-Bo Chen, Meng Hui Lim, and Mahesh Nanjundappa are kindly acknowledged for their analytical and pharmacokinetic work. We also appreciate Kah Fei Wan and his team for generating high-throughput antiviral activities. We are grateful to Caroline Rynn and her team for generating stability data in plasma, liver and intestinal S9. Ina Dix and Trixie Wagner are kindly acknowledged for their X-ray analysis of compound **17**. All authors were employees of Novartis at the time the work in this manuscript was performed.

